## Supplementary figures and images for "A dopamine-gated learning circuit underpins reproductive state-dependent odor preference in *Drosophila* females"

### Supplementary figures

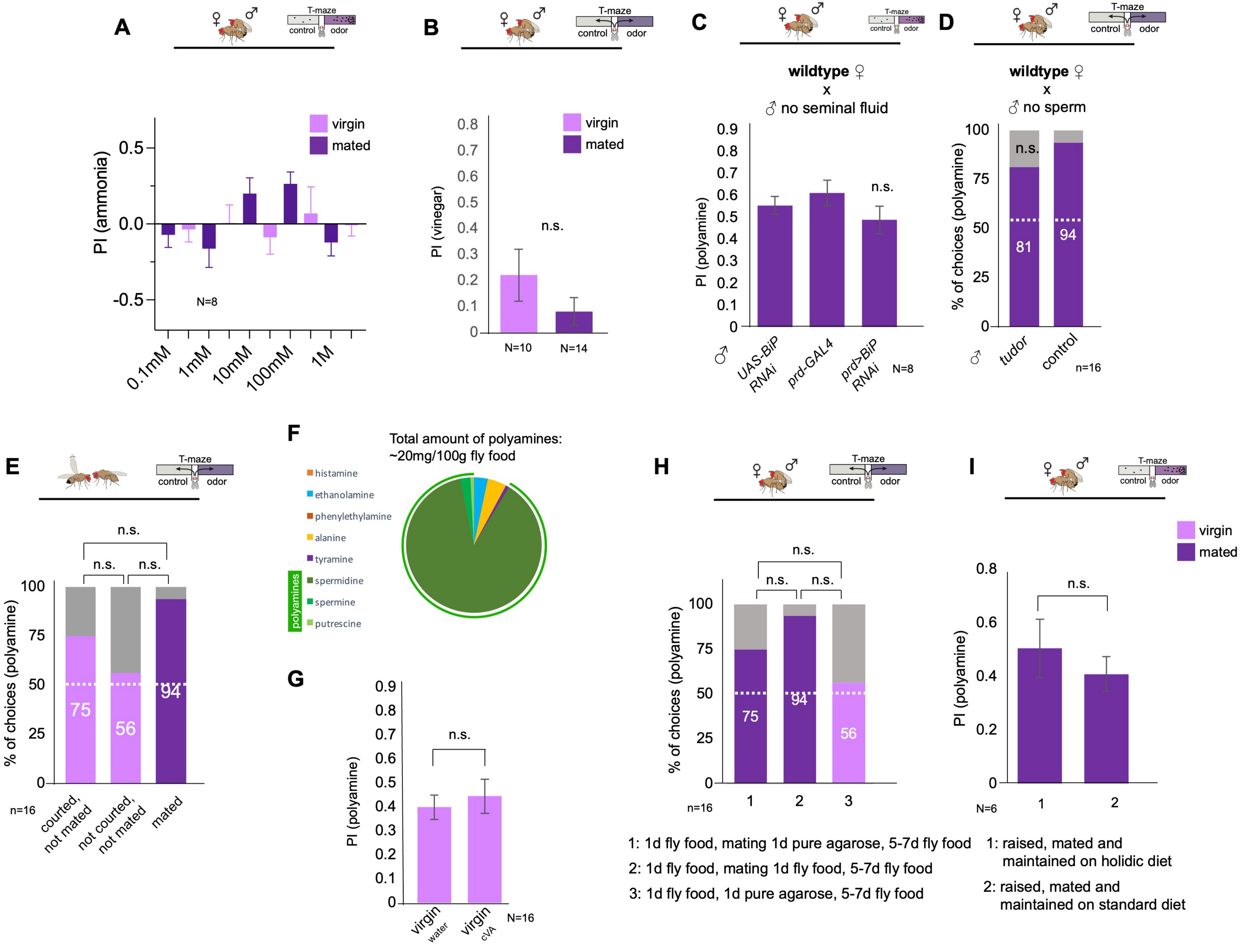

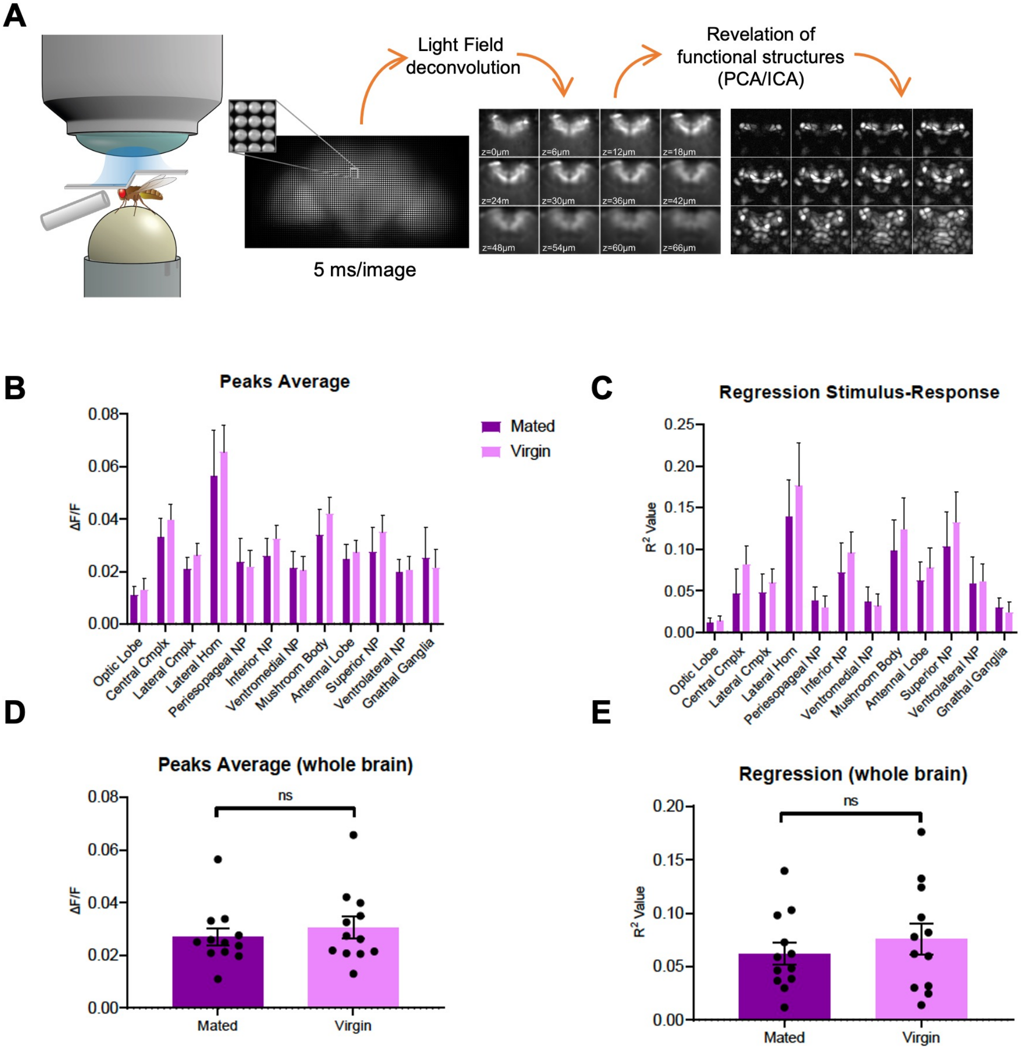

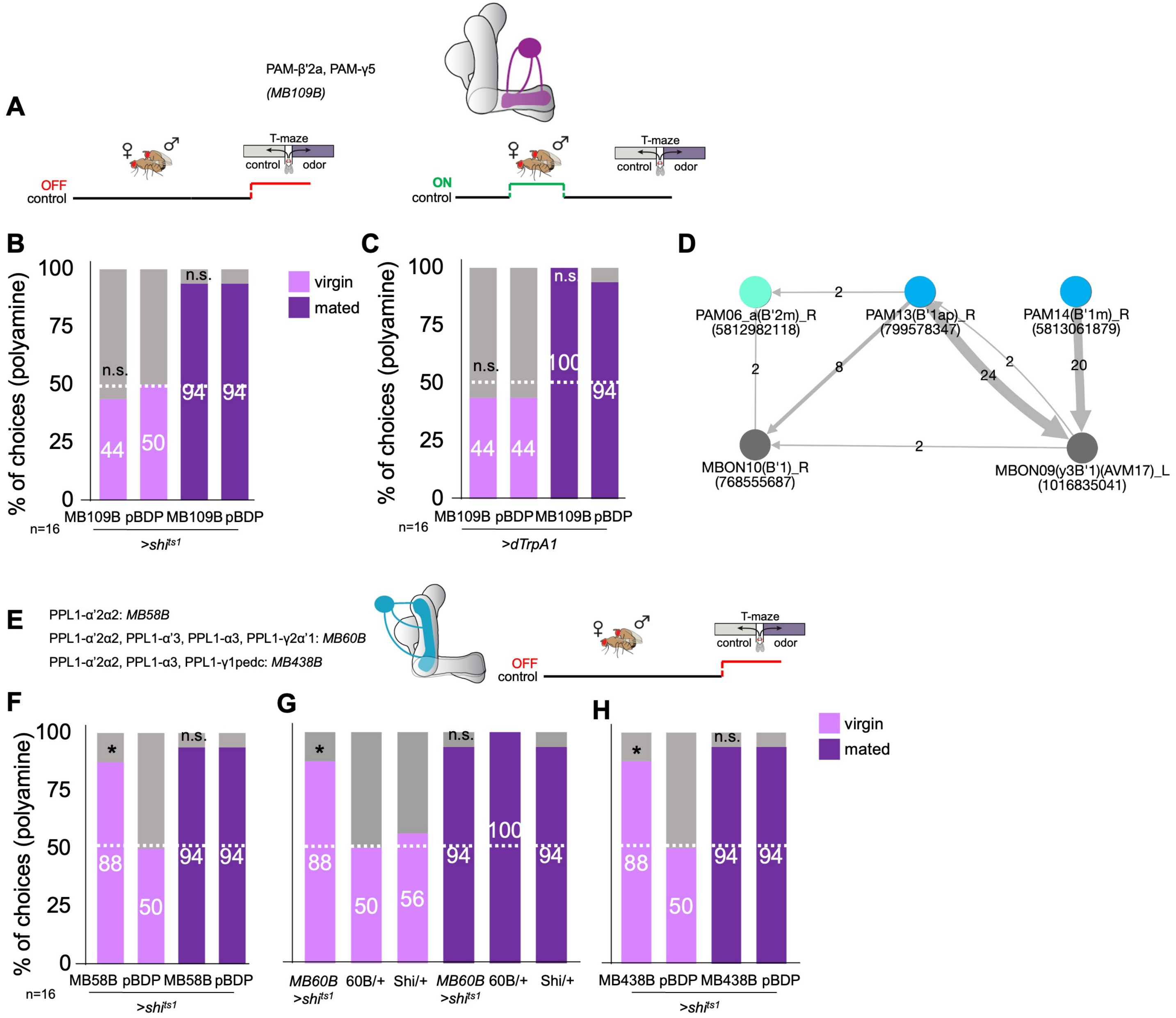

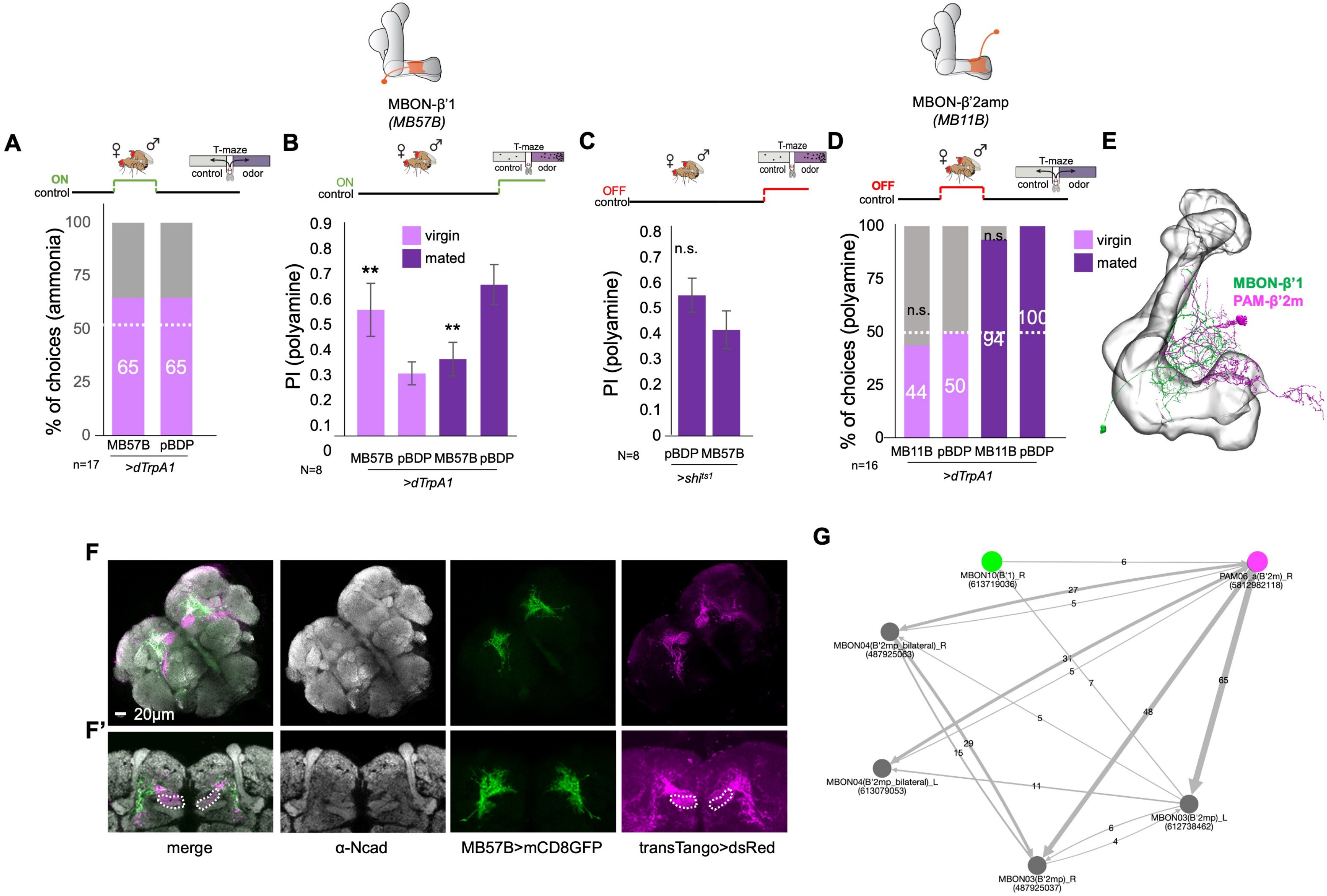

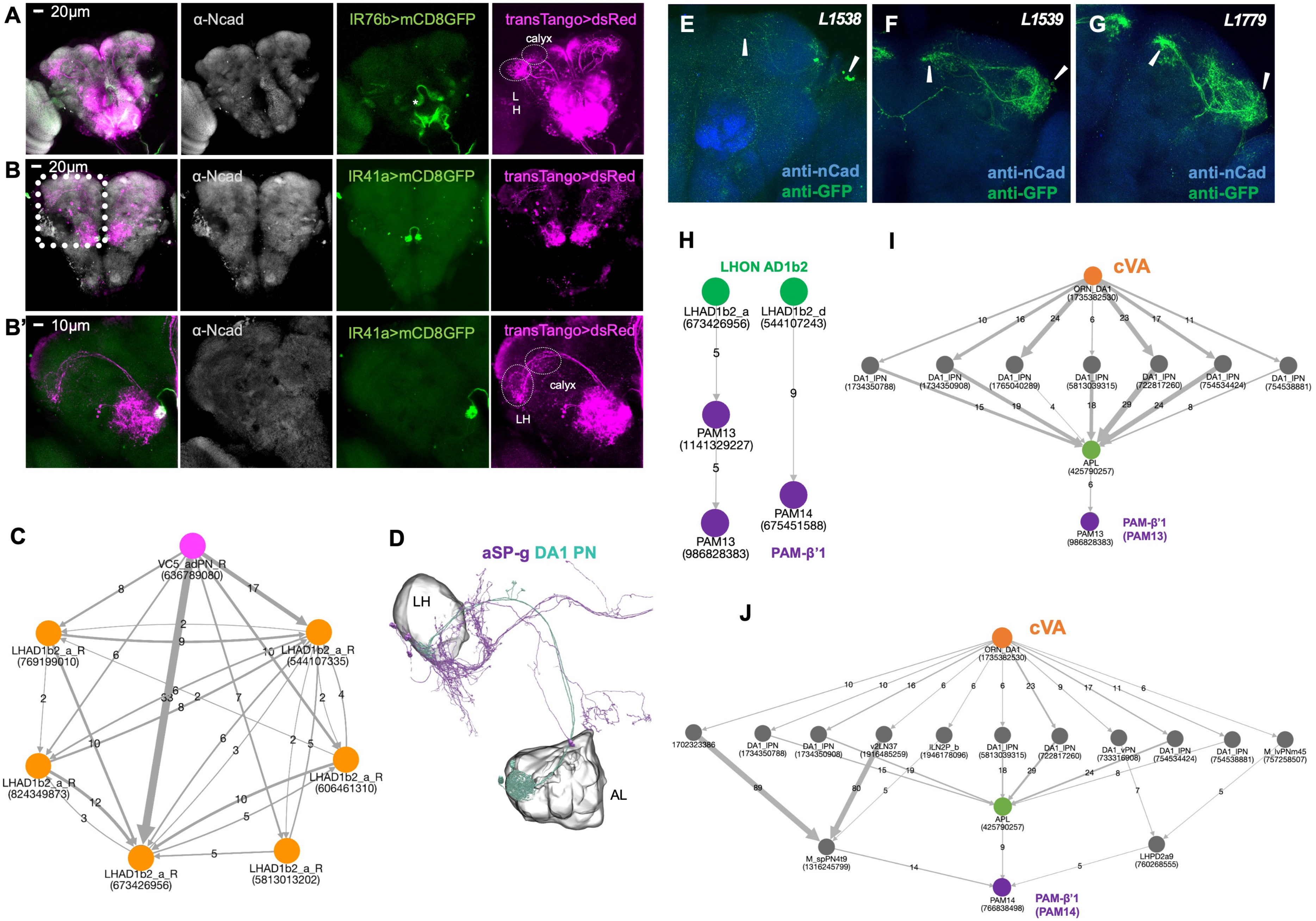

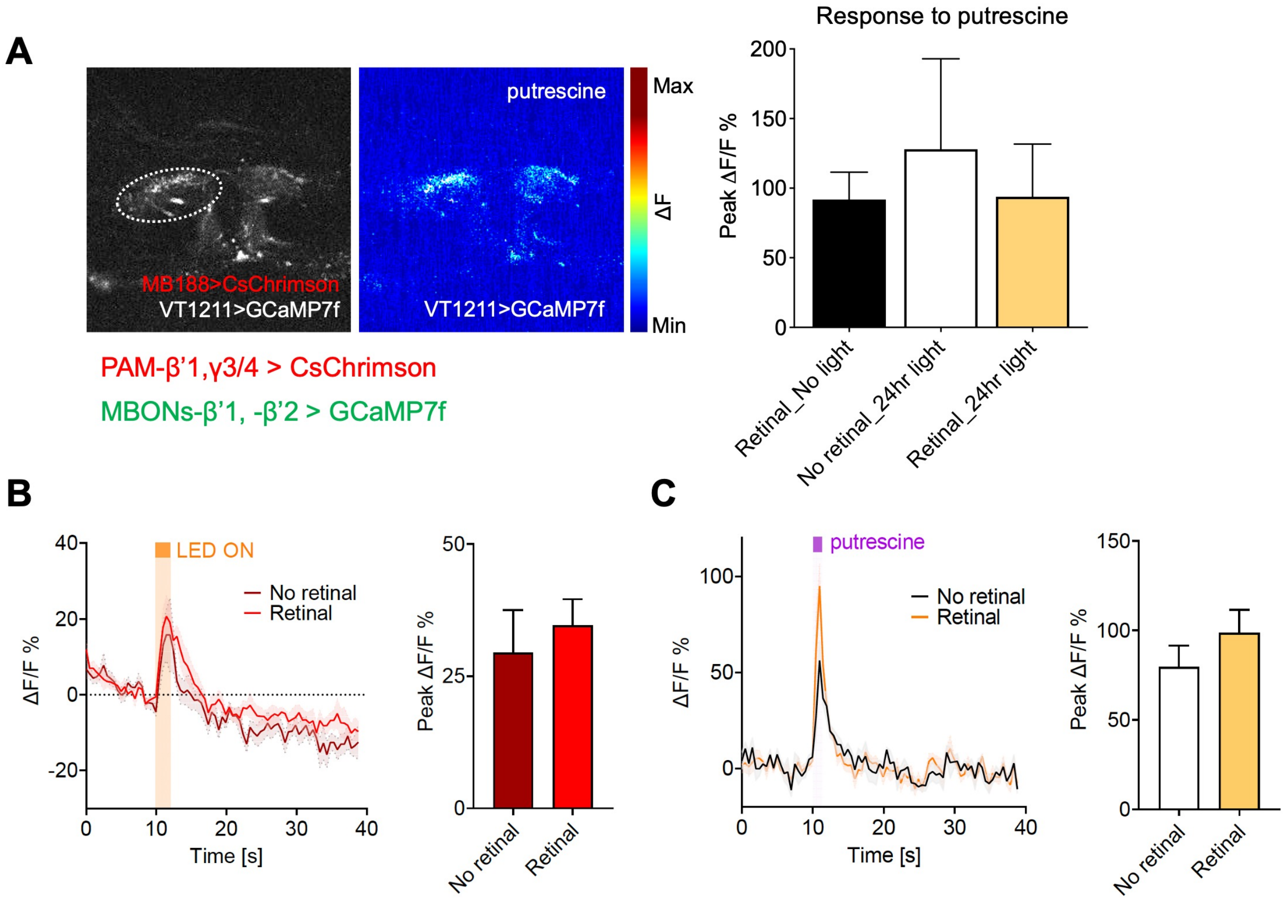

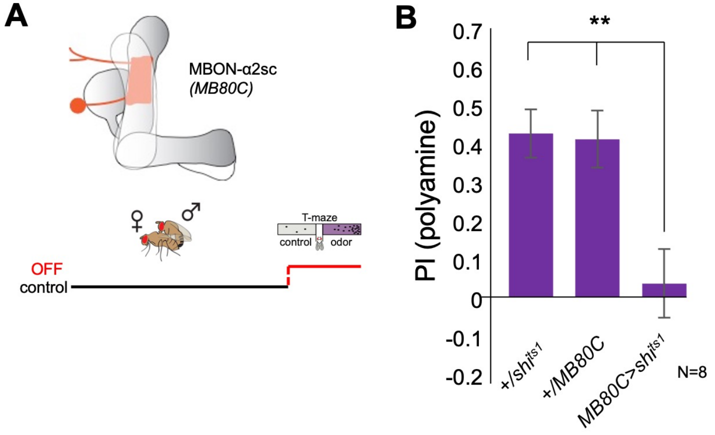
